# A nationwide genomic study of clinical *Klebsiella pneumoniae* in Norway 2001-2015: Introduction and spread of ESBL facilitated by CG15 and CG307

**DOI:** 10.1101/2021.07.16.452602

**Authors:** Aasmund Fostervold, Marit A.K. Hetland, Ragna Bakksjø, Eva Bernhoff, Kathryn E Holt, Ørjan Samuelsen, Gunnar Skov Simonsen, Arnfinn Sundsfjord, Kelly L Wyres, Iren H. Løhr, The Norwegian Study Group on *Klebsiella pneumoniae*

**Author notes:** **Collaborators**: The Norwegian Study Group on *Klebsiella pneumoniae:* Ståle Tofteland, Paul C. Lindemann, Nina Handal, Åshild M. Rødland, Aleksandra Jakovljev, Sandra Åsheim, Karianne W. Gammelsrud, Rolf A. Sandnes, Einar Weme, Angela Kümmel, Einar Nilsen, Belinda L. Lindstad, Anne Hollekim, Reidar Hjetland, Anne R. Oseid, Liv Jorunn Hafne. Corresponding author: Aasmund Fostervold, Avdeling for medisinsk mikrobiologi, Stavanger Universitetssykehus HF, Postboks 8100, 4068 Stavanger, NORWAY.

## Abstract

**Objective:** We have used the nationwide Norwegian surveillance program on resistant microbes in humans (NORM) to address longitudinal changes in the population structure *K. pneumoniae* isolates during 2001-15, encompassing the emergence and spread of ESBL-producing *Enterobacterales* (ESBL-E) in Norway.

**Material and methods:** Among blood (n= 6124) and urinary tract (n=5496) surveillance isolates from 2001-15, we used Illumina technology to whole genome sequence 201 ESBL-producing isolates from blood (n=130) and urine (n=71), and 667 non-ESBL isolates from blood. Complete genomes for four isolates were resolved with Oxford Nanopore sequencing.

**Results:** In a highly diverse collection*, Klebsiella variicola* ssp*. variicola* caused a quarter of *Klebsiella pneumoniae* species complex bacteraemias. ESBL-production was limited to *K. pneumoniae sensu stricto* (98.5 %). A diverse ESBL population of 57 clonal groups (CGs) were dominated by multidrug resistant CG307 (17%), CG15 (12%), CG70 (6%), CG258 (5%) and CG45 (5%) carrying *bla*_CTX-M-15_. Yersiniabactin was significantly more common in ESBL-positive (37.8%) compared to non-ESBL *K. pneumoniae sensu stricto* isolates (12.7%), indicating convergence of virulence and resistance determinants

Moreover, we found a significant lower prevalence of yersinabactin (3.0 %, 37.8 % and 17.3 %), IncFIB (58.7 %, 87.9 % and 79.4 %) and IncFII plasmid replicons (40.5 %, 82.8 % and 54.2%) in *K. variicola* ssp. *variicola* compared to ESBL- and non-ESBL *K. pneumoniae sensu stricto*, respectively.

**Conclusion:** The increase in Norwegian KpSC ESBLs during 2010-15 was driven by *bla*_CTX-M-15_ carrying CG307 and CG15. *K. variicola* ssp. *variicola* was a frequent cause of invasive KpSC infection, but rarely carried ESBL.

## Introduction

*Klebsiella pneumoniae* is an important human pathogen ^1^ and acknowledged as a key host for the spread of antimicrobial resistance (AMR)^2, 3^. The global spread of multidrug resistance (MDR) ^4^ *K. pneumoniae* is closely linked to the spread of extended-spectrum β-lactamases (ESBLs) and carbapenemases. This has been facilitated by successful clonal lineages or clonal groups (CGs) such as CG258, CG15 and CG307 ^5–7^, and horizontal gene transfer (HGT) ^2, 8^, fueled by antibiotic selection ^9^.

Whilst *K. pneumoniae* typically causes severe infections in vulnerable hospitalized patients ^1^, some hypervirulent (HV) clones cause community acquired invasive infections, often in healthy individuals ^10^. HV-clones cluster in CG23, CG65 and CG86, and harbour capsular loci K1 or K2, siderophores and other virulence factors supporting colonization, tissue invasion and immune evasion ^10, 11^. High-risk *K. pneumoniae* clones, categorized as either MDR or HV, rarely display both traits ^12^. However, in recent years, convergence of the two traits has been reported ^13^.

*K. pneumoniae* is a highly diverse species, and the term *K. pneumoniae* species complex (KpSC) has been introduced to encompass seven closely related taxa ^14^, of which *K. pneumoniae sensu stricto, Klebsiella variicola* and *Klebsiella quasipneumoniae* are the most frequently reported in human clinical samples ^15, 16^.

Most molecular epidemiological studies of KpSC have focused on outbreaks or isolates with particular characteristics such as AMR or virulence, most often with a cross-sectional study design. Thus, there is a need for longitudinal studies, including both resistant and susceptible isolates, to improve our understanding of the population dynamics in clinical KpSC isolates.

Here, we have used the Norwegian surveillance program on resistant microbes (NORM) during 2001-2015 to address the longitudinal dynamics of KpSC clinical isolates, dominant CGs and their associations with clinically important AMR- and virulence determinants.

The nationwide data show that the emergence of ESBL-producing KpSC clinical isolates in Norway has been dominated by MDR *bla*_CTX-M-15_ carrying *K. pneumoniae sensu stricto*, with CG15 and CG307 as major lineages. *K. variicola ssp. variicola* is a significant contributor to KpSC bacteraemias, but rarely carries ESBL genes in this geographical setting.

## Material and methods

### Bacterial isolates

NORM monitors AMR in *Klebsiella* spp. isolated from blood and urine isolates. Antimicrobial susceptibility is performed and interpreted according to EUCAST guidelines and breakpoints ^17^. Isolates with reduced susceptibility to cefotaxime and/or ceftazidime are categorized as ESBL or non-ESBL based on phenotypical ESBL-testing ^18^. Isolates are stored locally at −80°C at the participating laboratories ^18^.

All putative ESBL-producing KpSC blood (n=149) and urine isolates (n=91) from 2001-2015 registered in the NORM database were included in the study. For comparison, a subset of non-ESBL blood culture isolates (n=815) were included. To achieve a balanced sample for each year and to maintain a representative geographical and temporal distribution, consecutive entries were selected from each laboratory according to the following key: 2001, all isolates; 2005, every 2 out of 3; 2009, every 1 out of 2; and 2015, every 1 out of 3 (Figure S1).

Isolates registered as either *K. pneumoniae* or *Klebsiella* spp. were included. Species identification was confirmed by MALDI-TOF MS (MBT Compass Library DB-6903, Bruker Daltonik), and subsequently by whole genome sequencing. Only KpSC isolates were included for further analyses. Antimicrobial susceptibility profiles, laboratory, year and source of isolation were retrieved from the NORM database. For isolates with discordant ESBL geno- and phenotype, the phenotype was confirmed using the combined disc method (Becton Dickinson, New Jersey, USA) ^19^. Colistin MIC was determined by broth microdilution using Sensititre FRCOL plates (Thermo Fisher Scientific, East Grinstead, UK) according to manufacturer’s instructions.

### Whole genome sequencing and *in silico* analysis

See supplementary methods for details. Paired-end reads (300 or 125 bp) were generated for all isolates using Illumina MiSeq and Illumina HiSeq platforms, respectively. Selected isolates were also long-read sequenced on a MinION Mk1B device (Oxford Nanopore Technologies). Unicycler v0.4.8 ^20^ was used for all assembly.

Kleborate v2.0.4 ^21^ was used to identify species, sequence type (ST), virulence loci and AMR genes (CARD database v3.0.8 ^22^). Kaptive ^23^ was used to identify capsule (K) biosynthesis loci reporting calls with confidence level “Good” or higher.

Read-sets from putative ESBLs with no definite ESBL gene were investigated using SRST2 v.2.0.4 ^24^ with CARD database v3.0.8, as reads containing ESBL sequences may have been discarded during assembly, and/or assembled across multiple contigs. Plasmid replicons were identified with SRST2 v0.2.0 using the PlasmidFinder database version 2021-01-13 ^25^.

A core chromosomal single-nucleotide variant (SNV) alignment of all genomes was generated by mapping short-reads to the chromosome of the ST23 reference genome NTUH-K2044 (Genbank accession NC_01273.1) using RedDog V1beta.11 ^26^. A maximum likelihood (ML) phylogeny was inferred from the resulting alignment using FastTree v2.1.10 ^27^.

CGs were defined from the phylogeny using patristic distance cut-off of 0.04 ^13^ and named according to the dominant ST within each CG. However, CGs dominated by ST14 and ST340 were named CG15 and CG258, as these are more commonly known^2, 28^. An alignment of all CG307 genomes was generated with RedDog V1beta.11, using the closed ST307 genome (Genbank accession CP073627) as the mapping reference, and subsequently passed to RAxML v8.2.10 ^29^ to infer a CG307 clone-specific ML phylogeny.

### Data availability

The 868 KpSC short-read and three long-read sequence files have been deposited in the European Nucleotide Archive under BioProject PRJEB27256 (Table S1). The four hybrid-assembled completed genomes have been deposited in GenBank (Table S1) under accession numbers CP073791-CP073796, CP073627-CP073629, CP073783-CP073787 and CP073788-CP073790.

### Definitions

MDR was defined as phenotypic resistance to agents in three antimicrobial classes ^4^. HV was defined as either a) the presence of *rmpA* or *rmpA2;* and/or b) the presence of aerobactin (*iuc*) and salmochelin (*iro*) ^30^. ESBL isolates were defined as either having known ESBL genes (i.e. *bla*_CTX-M_, *bla*_SHV-2_, *bla*_SHV-5_, *bla*_SHV-12_, *bla*_SHV-18_ and *bla*_SHV-24_), or in absence of known ESBL genes, a confirmed ESBL phenotype. Isolates with plasmid-mediated AmpC genes only or carbapenemase-encoding genes (regardless of ESBL gene presence) were excluded from further analysis.

### Data handling and statistical analysis

Data analysis and statistics was done using R version 4.0.2 (2020-06-22) ^31^. Distribution differences were calculated with Fisher exact test, with Benjamini-Hochberg correction for multiple testing when necessary. p < 0.05 was considered statistically significant.

### Ethics

The study was approved by the Regional Committee for Medical and Health Research Ethics (Reference: 2017/1185-3).

## Results

We received 954/1,055 (90.4%) of requested isolates of which 223 putative ESBL-producing isolates (blood, n=144; urine, n=79) and 667 non-ESBL blood isolates were confirmed as KpSC by MALDI-TOF MS and WGS (Figure S1). Known ESBL genes were detected in 192/223 (86%) putative ESBL isolates. ESBL phenotype was confirmed in nine additional isolates, resulting in an ESBL group consisting of 201 isolates (blood, n=130; urine, n=71). Six isolates with carbapenemase genes and one isolate with a plasmid-mediated AmpC gene only were excluded (Figure S2). The dataset can be explored at https://microreact.org/project/4dBcaZsZmKoAzvatzPaGds.

### Phylogenetic diversity in ESBL and non-ESBL KpSC populations

The species distributions in the ESBL and non-ESBL groups were different (p<0.0001) (Table 1). The ESBL group consisted of 98.5% *K. pneumoniae sensu stricto*. In contrast, the non-ESBL group isolates comprised *K. pneumoniae sensu stricto* (69.1%), *K. variicola* ssp. *variicola* (24.5%), and *K. quasipneumoniae* ssp. *similipneumoniae* (3.3%) and *K. quasipneumoniae* ssp. *quasipneumoniae* (3.1%). *K. variicola* ssp. *tropica, K. africana* or *K. quasivariicola* were not detected (Table 1, Figure 1).

**Table 1.** ESBL and non-ESBL groups: species distribution, clonal groups (CGs), and sequence types (STs) numbers.

| Species identification | ESBL isolates (%) | ESBL (CGs /STs) | non-ESBL isolates (%) | non-ESBL (CGs/STs) | p<sup>1</sup> |
| --- | --- | --- | --- | --- | --- |
| *K. pneumoniae sensu stricto* | 198 (98.5) | 54/70 | 461 (69.1) | 136/222 |  |
| *K. variicola ssp. variicola* | 1 (0.5) | 1/1 | 163 (24.5) | 80/115 |  |
| *K. quasipneumoniae ssp. similipneumoniae* | 2 (1) | 2/2 | 22 (3.3) | 20/20 |  |
| *K. quasipneumoniae ssp. quasipneumoniae* | - | - | 21 (3.1) | 10/20 |  |
| *K. pneumoniae* species complex | 201 | 57/73 | 667 | 246/377 | <0.0001 |

<sup>1</sup>Fisher exact test for difference in species distributions between groups

**Figure 1.**
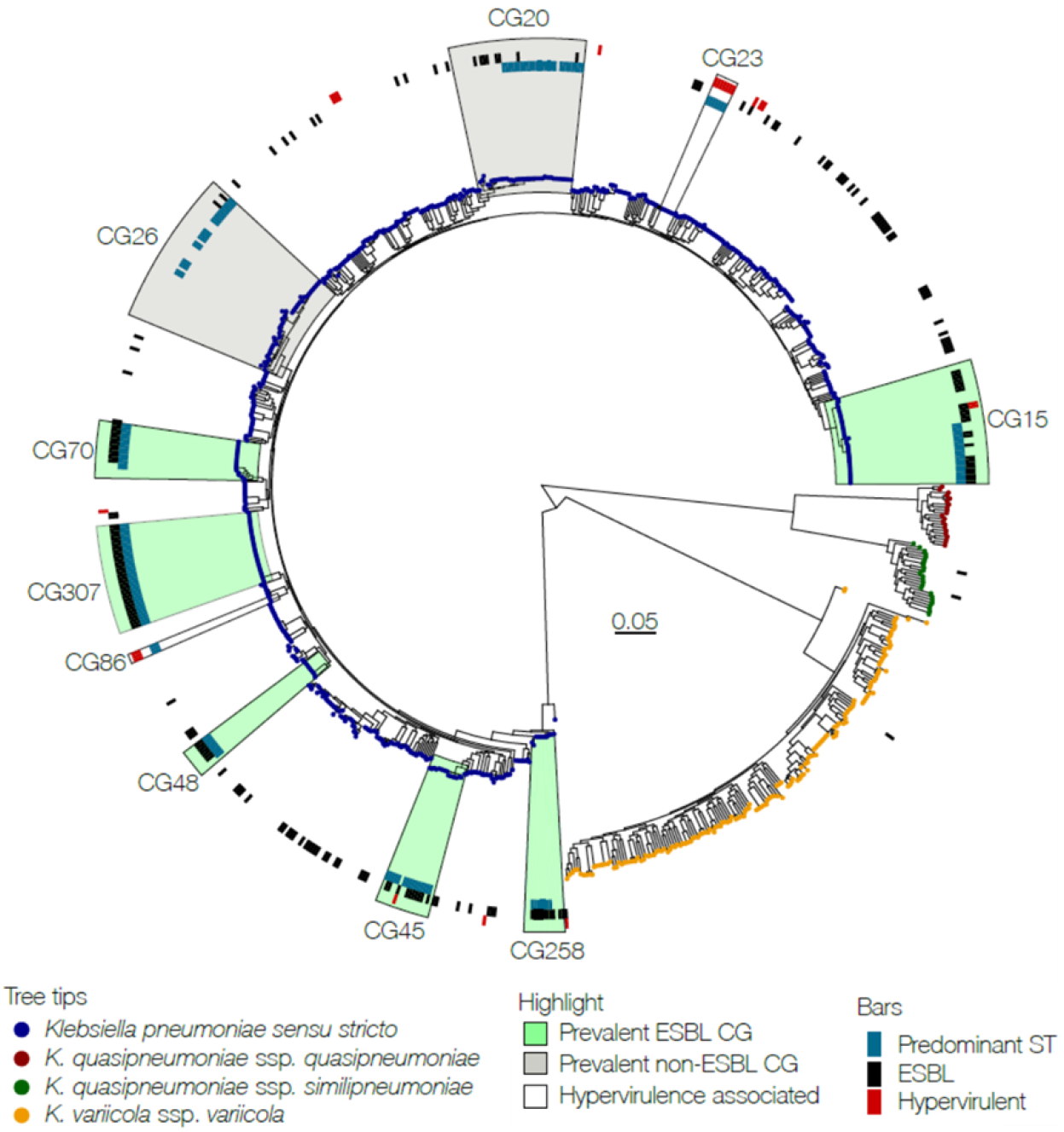
Maximum likelihood tree of the 868 *K. pneumoniae* species complex genomes. Tips are colored by *Klebsiella* species. CGs each representing >5% of isolates either in the ESBL or the non-ESBL group, are highlighted in green and grey, respectively. CGs commonly associated with hypervirulence are highlighted without color. The highlighted areas include all genomes on the most recent common ancestor node of the CG indicated. The circles from inner to outer show genomes with the predominant sequence type (ST, blue bars) within the CG, presence of ESBL-encoding genes (black bars) and isolates meeting the study definition of hypervirulence (red bars).

The 868 KpSC isolates were phylogenetically highly diverse with a total of 413 different STs assigned to 261 CGs (Table S1). The Simpsons diversity indices for STs were 0.95 for the ESBL group and 0.99 for the non-ESBL group, respectively.

The ESBL group (n=201) consisted of 73 STs and 57 CGs (mean number of isolates per CG 3.53, range 1-34). CG307 was the most prevalent (16.9%, n=34; ST307) followed by CG15 (12.4%, n=15; ST14, ST15 and ST627), CG70 (6.5%, n=13; ST70), CG258 (5.0%, n=10; ST11, ST340 and ST437) and CG45 (5.0%, n=10; ST45 and ST2954). The remaining CGs represented less than 5% of isolates each (Figure 1).

Among the 667 non-ESBL blood culture isolates there were 377 STs and 246 CGs (mean number of isolates per CG 2.71, range 1-42). CG26 (6.3%, n=42; 11 STs) and CG20 (5.2%, n=35; 6 STs) were the most prevalent. In the remaining 244 CGs (88.5%, n=590) each CG represented less than 5% of the isolates. Ten (1.5%) Hv-associated isolates, CG23 (n=7) and CG86 (n=3), were observed. CG307 was absent in the non-ESBL group, while the other major ESBL CGs were present in low numbers; CG15 (2.1%, n=14), CG70 (0.7%, n=7), CG45 (1%, n=7) and CG258 (0.4%, n=3). In total, 33 of 57 CGs (57.9 %) in the ESBL group were present in the non-ESBL group.

### Temporal trends

The globally successful ESBL CG15 was first observed in this study in 2003, in a urine specimen, becoming the most prevalent CG in blood culture samples between 2009 and 2012. From 2012, the increase in ESBLs was associated with the emergence of CG307. Urine isolates exhibited greater diversity of CGs compared to blood, but all prevalent urine CGs were also represented in blood cultures isolates, albeit several in low numbers (Figure 2). There were no apparent CG-trends in the non-ESBL group.

**Figure 2.**
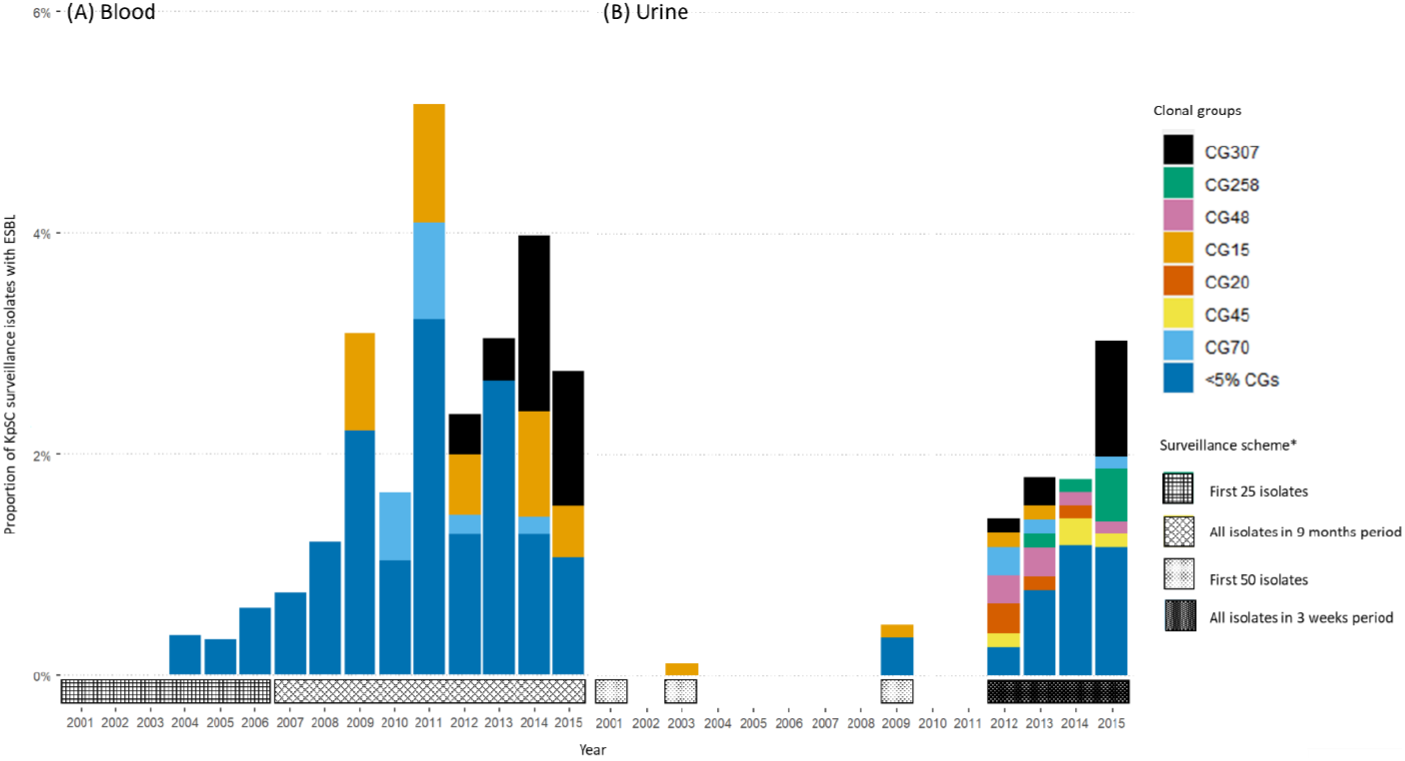
Temporal distribution of *Klebsiella pneumoniae* species complex (KpSC) ESBL clonal groups (CGs) in blood (A) and urine (B) as proportion of the surveillance for each year. Distribution of CGs in blood and urine separated by most prevalent CGs and other CGs. *The surveillance starts in January each year.

CG307 was first observed in 2012 in two of the six NORM surveillance regions, and present in five regions by 2014 (Figure 3A), representing 44.4% of blood- and 34.6% of urine ESBL-isolates in 2015. CG70, CG258 and CG45 emerged in the same period, but did not expand to the same degree. Chromosomal single-nucleotide variants (SNVs), temporal and geographical distribution of the most prevalent CGs are shown in Table S5. A core genome phylogenetic analysis of CG307 indicated independent occurrences of isolates in 2012-2013, while isolates from 2014-2015 seem to represent two clonal expansions (red and blue boxes figure 3B). This is supported by the Bayesian phylodynamic analysis of global CG307 (including 30 of the genomes reported here), adapted from Wyres *et al.* ^6^, showing that the most recent common ancestor for the two proposed clonal expansions date back to 2009 (Figure S3).

**Figure 3.**
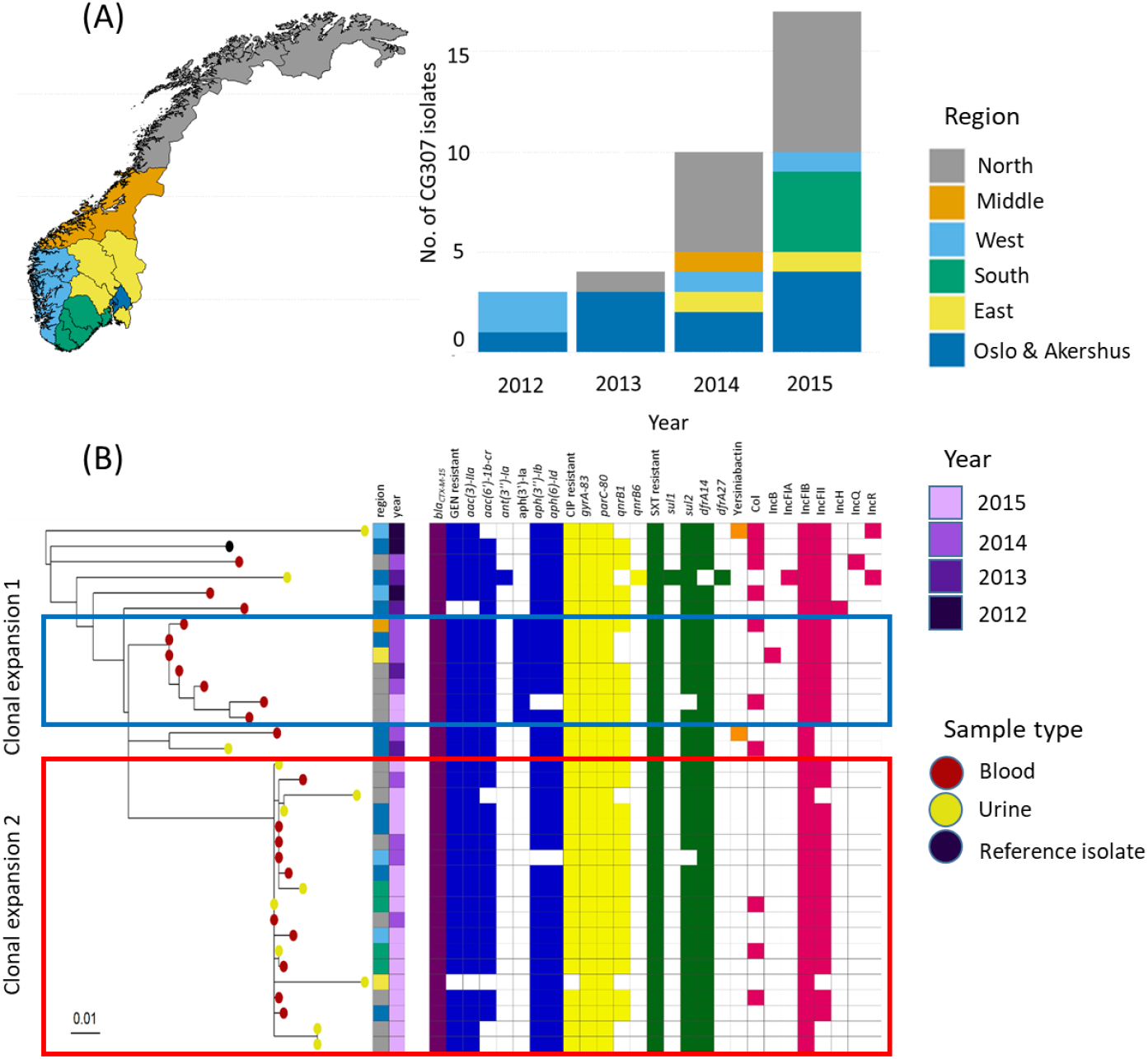
Epidemiology and phylogeny of CG307. **(A)** Map of Norway with surveillance regions as defined by the Norwegian surveillance program on resistant microbes (NORM), with bar plot showing number of CG307 isolates per region per year. **(B)** CG307 core genome phylogeny with metadata, distribution of AMR determinants, virulence determinants and replicon families. Red and blue boxes mark proposed clonal expansions.

### Antimicrobial resistance – phenotype and genotype

All ESBL group isolates had an ESBL phenotype. Overall, *bla*_CTX-M-15_ was the most prevalent genotype (n=146; 72.6%), more dominant in high prevalence CGs (93.5%) compared to other CGs (55%, p <0.001). The remaining ESBL-genotypes comprised a diversity of *bla*_CTX-M_ (n=22; 10.9 %) or *bla*_SHV_ (n=22; 10.9 %) (Figure 4). Reduced susceptibility to meropenem were found in two ESBL isolates (CG258) and one non-ESBL isolate (CG515), all without carbapenemase genes.

**Figure 4.**
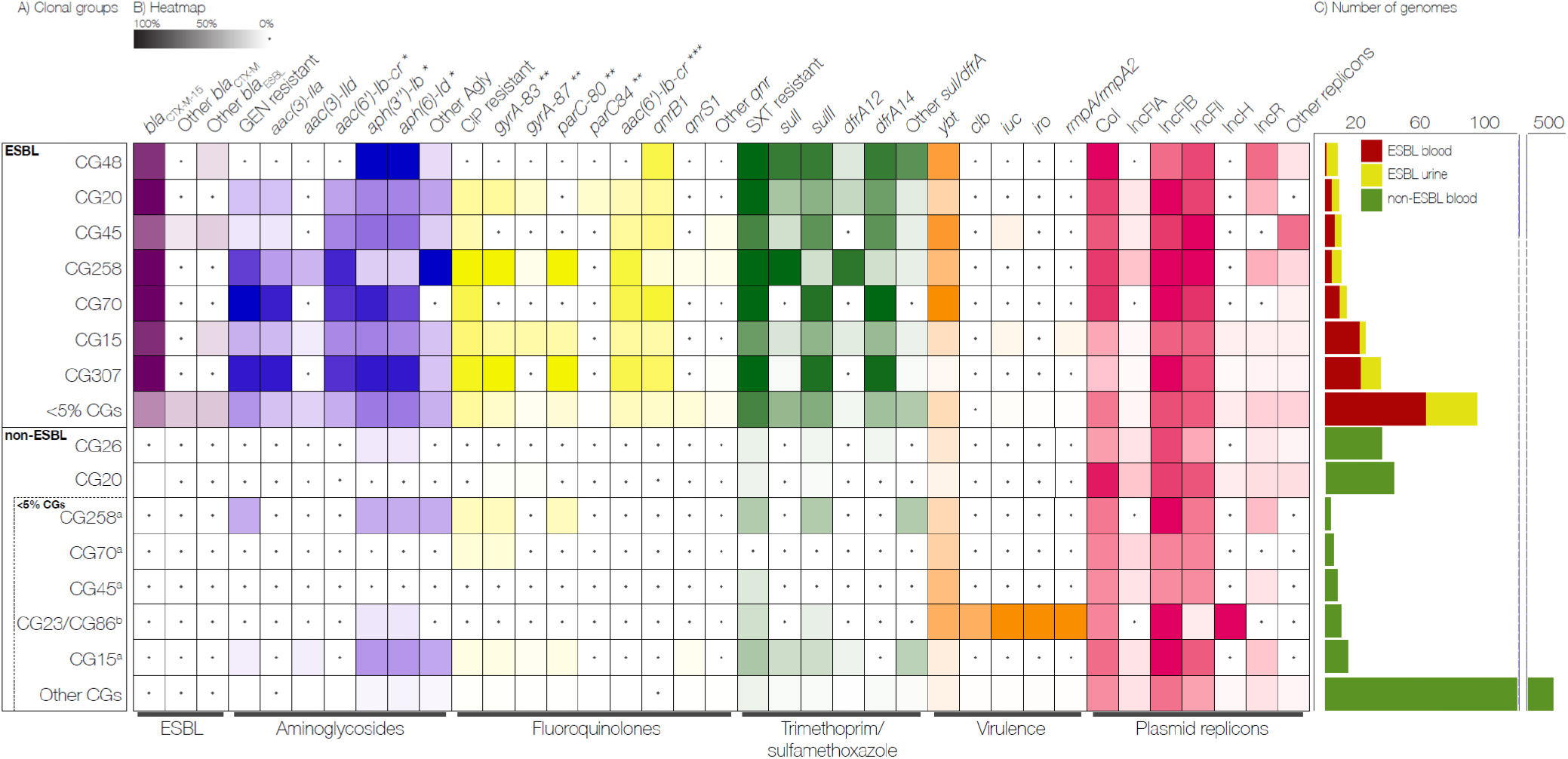
Distribution of resistance determinants, virulence factors and plasmid replicons among clonal groups (CGs). **A)** Distribution of CGs in the ESBL and non-ESBL groups, separated by most prevalent CGs (each representing >=5% of isolates in group) and other CGs. In the non-ESBL group, <5% CGs which were the **a)** most prevalent CGs from the ESBL group and the **b)** hypervirulence-associated CG23 and CG86 are shown separately. **B)** The intensity of the box shading indicates the percentage of genomes harboured determinant. White shading with a black dot indicates that there are no determinants present. For each antibiotic class, the presence of resistant phenotype and resistance determinants are indicated. **GEN** = gentamicin, **CIP** = ciprofloxacin, **SXT** = trimethoprim-sulfamethoxazole, **Agly** = acquired aminoglycoside resistance genes, **\*** Does not confer resistance to gentamicin, **\*\*** Chromosomal mutation position, **\*\*\*** May reduce susceptibility to both aminoglycosides and fluoroquinolones, **C)** Total number of genomes in the ESBL and non-ESBL groups.

In the ESBL group, reduced susceptibility to gentamicin, ciprofloxacin and trimethoprim-sulfamethoxazole was found in 54.7% (n=110), 72.1% (n= 145) and 88.1% (n=177) of the isolates, with at least one corresponding AMR determinant found in 95.5 % (n =105), 97.8% (n=142) and 96.6 % (n= 171) of the isolates, respectively.

Prevalent AMR-determinants are shown in Figure 4. Of note, *armA* (n=1) and *rmtG* (n=1) encoding 16S-rRNA methylases were rare, while *aac(6’)-Ib-cr* which may reduce susceptibility to both aminoglycosides and quinolones was found in 48.3 % (n= 97) of the ESBL isolates. Interestingly, *aac(6’)-Ib-cr* was frequently found along with other determinants, in particular in 82.5 % (n=66) and 84.0 % (n=68) of isolates carrying *aac(3)-IIa* and *qnrB1*, respectively.

Colistin resistance determinants were rare. Three ESBL isolates carried *mcr-9.1* (n=2) or a truncated chromosomal gene *mgrB* (n=1) where only the isolate with truncated *mgrB* showed an elevated MIC of 16 mg/L for colistin.

In the non-ESBL group, reduced susceptibility to gentamicin, ciprofloxacin and trimethoprim-sulfamethoxazole was observed in 0.9% (n=5), 2.5% (n=17) and 9.4% (n=63) of the isolates, with at least one corresponding AMR determinant found in 80.0 % (n =4), 58.8% (n=10) and 60.3 % (n= 38), respectively. Prevalent AMR-determinants are shown in Figure 4.

MDR was frequent in the ESBL group, (70.1 %, n= 141) compared to the non-ESBL group (0.3 %, n=2), (p< 0.001). Notably, 83.0 % (n=117) of MDR ESBL isolates were carrying *bla*_CTX-M-15_. Only 11.2% (n=18) of ESBL isolates were susceptible to all three of gentamicin, ciprofloxacin and trimethoprim-sulfamethoxazole.

### Diversity in capsule loci and virulence determinants

Capsule loci (KLs) were identified in 73.6 % (n=148, 38 KLs) of the ESBL isolates and 54.3 % (n=362, 87 KLs) of the non-ESBL isolates. KL102 (13.4 %, n=27) was the most prevalent KL among the ESBL isolates, mainly associated with CG307 (n=25). The HV-associated KL1 and KL2 were rare, detected in 5.5 % of ESBL (KL2, n=11) and 3.7% of non-ESBL (KL1, n=11; KL2, n=14) isolates, respectively (Table S2).

Eighteen isolates (2.1 %), all *K. pneumoniae sensu stricto*, met the HV definition, where seven of 15 non-ESBL HV isolates belonged to CG23 (Table S3). Long-read sequencing of one CG133 (ST420) (Genbank accession CP073783-CP073787) ESBL isolate showed *bla*_CTX-M-15_ to be situated on an IncFII plasmid without any of the virulence loci. However, two ST15 ESBL isolates (BioSample accession SAMEA5063299, SAMEA5063230), harboured the iuc and rmpA2 virulence loci, as well as blaCTX-M-15 on the same mosaic plasmid, as described by Lam et al. ^32^.

The distribution of virulence determinants is shown in Figure 4 and Table S3. Yersiniabactin was the most prevalent acquired siderophore in *K. pneumoniae sensu stricto* isolates, and more dominant in ESBL isolates (37.8%, n=75) compared to non-ESBL isolates (17.3 %, n=80) (p < 0.001). Notably, among *K. pneumoniae sensu stricto* non-ESBL isolates, there was higher prevalence of yersiniabactin in CGs that were also found in the ESBL group compared to other CGs (29.4 % and 11.0% respectively, p<0.0009). Only 5/163 (3.1 %) non-ESBL *K. variicola spp. variicola* had yersiniabactin. While no ESBL isolates had the genotoxin colibactin, it was present in 11 non-ESBL isolates (CG23, n=7; CG133, n=2; CG417; CG643).

### Plasmid replicon patterns in ESBL and non-ESBL KpSC populations

Fifteen plasmid replicon families were identified in the ESBL group. IncFIB (87.9%, n=176) and IncFII (82.8%, n= 164) were the most common. In the non-ESBL group, fourteen plasmid replicon families dominated by IncFIB (72.0%, n=480), IncFII (48.7%, n=325), IncFIA (15.0%, n= 100), and IncR (14.5%, n= 97) were identified. Twelve plasmid replicon families were found in both groups (Table S4).

Interestingly, IncFIB and IncFII were more abundant in *K. pneumoniae sensu stricto* in the ESBL group compared to the non-ESBL group (p<0.001 and p <0.001, respectively). Additionally, in the non-ESBL group IncFIB and IncFII were significantly more common in *K. pneumoniae sensu stricto* compared to the other species (p<0.001 and p<0.03, respectively, Table 2).

**Table 2.** Distribution of prevalent replicon types in *Klebsiella pneumoniae sensu stricto* and *K. variicola* ssp*. variicola*.

|  | ESBL | Non-ESBL |  | p-value* |  |
| --- | --- | --- | --- | --- | --- |
| Replicon type | A.<br>*K. pneumoniae sensu stricto*<br>(n=198) | B.<br>*K. pneumoniae sensu stricto*<br>(n=461) | C.<br>*K. variicola ssp. variicola*<br>(n=163) | A vs B | B vs C |
| Col | 63.1% (125) | 63.3% (292) | 47.9% (78) | NS | 0.006 |
| IncFIA | 6.1% (12) | 14.3% (66) | 17.8% (29) | 0.004 | NS |
| IncFIB | 87.9% (174) | 79.4% (366) | 58.3% (95) | 0.001 | 0.001 |
| IncFII | 82.8% (164) | 54.2% (250) | 40.5% (66) | 0.001 | 0.03 |
| IncR | 18.2% (36) | 14.1% (65) | 12.3% (20) | NS | NS |

\*Fisher exact test with Benjamini & Hochberg correction for multiple testing. NS – not significant.

## Discussion

We have used the nationwide Norwegian AMR surveillance framework to perform a population structure analysis of all ESBL producing KpSC blood and urine isolates as well as a representative collection of non-EBSL blood isolates during 2001-15 for comparison. The combined use of WGS and national registry data, allowed the analysis of temporal and geographical trends in the species distribution, phylogeny, AMR and virulence determinant content in KpSC clinical isolates during a period when ESBL-producing *Enterobacterales* gained foothold in Norway.

Firstly, we noted a significant difference in species distribution between the ESBL and non-ESBL group. While the ESBL-group was essentially dominated by *K. pneumoniae sensu stricto, K. variicola* ssp. *variicola* accounted for 24.5% of the non-ESBL group. This is in line with findings in ESBL-producing or MDR KpSC-strain collections from the USA ^33^ and the British Isles ^33, 34^, dominated by *K. pneumoniae sensu stricto*. Two studies (Sweden 2007-09, single centre ^16^ and Japan 2014-17, two-center) of consecutive blood KpSC isolates, both with low prevalence of ESBL, showed similar species distributions compared to our results.

As gastrointestinal colonization is considered the primary source of the majority of KpSC bloodstream infections ^35, 36^, we expect that the observed proportion of *K. variicola* ssp. *variicola* among blood isolates reflects the ratio of gut colonization in the patient population. This is supported by a recent Norwegian study where 16.3% of 2,975 healthy adults had KpSC in faecal screening samples, of which 28% were *K. variicola spp. variicola* ^37^. While *K. pneumoniae sensu stricto* is the predominant KpSC species reported in other gut carriage studies, carriage rates in the range of 10-20% of *K. variicola* have been shown for intensive care patients ^35^ and pregnant women in low-income countries ^30^. As a frequent gut resident, one could expect *K. variicola spp. variicola* to acquire ESBL-encoding plasmids and genes *in vivo*. To our knowledge there are no experimental studies supporting any mechanisms explaining the low abundance of ESBL in *K. variicola spp. variicola* compared to *K. pneumoniae sensu stricto*.

ESBL rates increased in clinical KpSC isolates during the study period, from 0% in 2001 to 3.1% in 2015 ^18^. Our temporal data show an increasing predominance of *bla*_CTX-M-15_ accompanied by an increasing co-resistance to other clinical important antibiotics. An overall increase in ESBL and MDR rates in clinical isolates of KpSC was observed in other European countries in the same period ^38^. AMR determinants, as well as phenotypic resistance against gentamicin, ciprofloxacin and trimethoprim-sulfamethoxazole, were rare among non-ESBL isolates in our study. This observation strongly suggest that the overall increase in Norwegian MDR KpSC is driven by the expansion of ESBL-producing *K. pneumoniae sensu stricto*, in particular *bla*_CTX-M-15_ carrying CGs, such as CG15 and CG307.

Our KpSC strain collection is characterized by a large clonal diversity, both in the ESBL and non-ESBL groups, throughout the study period. The most striking shift in the temporal data, in addition to increasing ESBL rates, is the introduction and subsequent spread of CG307 since 2012. The CG307 phylogenetic analyses based on our dataset reveal nationwide expansion of this clone, which seems to be closely related to international isolates as previously shown by Wyres *et al.* ^6^. Notably, we have not observed non-ESBL CG307. In contrast, clinical isolates of CG15, a frequent carrier of ESBL, were also present without ESBL.

The observed clonal diversity within the ESBL group and the increasing abundance of CG307 and CG15, is in line with results reported by Moradigaravand *et al.* in the British isles ^34^ and Long *et al.* in the USA ^33^ during 2001-2011 and 2011-2015, respectively. In contrast to their findings, CG258 was less frequently detected in our study. CG307 is still playing an important role in the dissemination of ESBL KpSC in Norway, as confirmed by WGS of blood culture ESBL isolates reported to NORM in 2019, where CG307 (29.2%) remains the dominant CG ^18^. While the prevalence of carbapenemase-producing KpSC in Norway is still low ^18^, the establishment of ESBL CG307 is a cause for concern, as this clone has shown to be a well prepared host for carbapenemase genes in other settings ^7^.

Our results suggest that HV KpSC, including CG23 and CG86, are rare in clinical isolates in Norway. This is also in line with the recent population structure analysis of Norwegian KpSC faecal carrier isolates ^37^. Yersiniabactin, usually transferred by integrative conjugative elements (ICEs) ^39^, is the most prevalent virulence-associated gene, mainly found in *K. pneumoniae sensu stricto* isolates. Notably, we observed a significantly higher prevalence of yersiniabactin in ESBL (37.8%) compared to non-ESBL (17.3 %) *K. pneumoniae sensu stricto* isolates, in contrast to previously published results ^39^. Other virulence determinants were uncommon and played no dominant role in the examined KpSC population. However, as previously reported by Lam *et al.*, virulence-encoding genes were found in convergence with *bla*_CTX-M-15_ on mosaic plasmids in two ST15 isolates ^32^.

Our data demonstrate that the replicon families IncFIB and IncFII, frequently associated with ESBL genes ^40^, were common both in ESBL (87.9% and 82.8 %) and non-ESBL (79.7% and 54.2 %) *K. pneumoniae sensu stricto* isolates. These replicons were also present in *K. variicola* ssp. *variicola* (58.3% and 40.5 %), but significantly less prevalent compared to non-ESBL *K. pneumoniae sensu stricto*.

Importantly, our data support the notion that there seems to be a higher propensity for certain *K. pneumoniae sensu stricto* clonal groups to acquire mobile genetic elements, represented by ESBL-encoding plasmids and yersinabactin-linked ICEs, compared to *K. variicola* ssp. *variicola*. This is concordant with Wyres et al. ^13^ showing some KpSC clones to be generally better at acquiring genetic material via horizontal gene transfer than others. The observations need further investigations including experimental studies of underlying mechanisms.

The strength of this study lies in the use of the comprehensive unselected national surveillance data collected over a 15-year period, encompassing the introduction of ESBL, and by using WGS gaining detailed insight into genomic epidemiological features. As we opted for temporal and geographical diversity, we have not done a randomized selection of non-ESBL isolates, which may have introduced a bias in estimating the prevalence of significant CGs or genetic determinants in recent years. The lack of urine ESBL isolates in the periods 2004-2008 and 2010-2011 may also conceal the early appearance of significant CGs.

In conclusion, the increase of ESBL and clinically relevant co-resistance in *K. pneumoniae sensu stricto* in Norway during the study period is closely linked to *bla*_CTX-M-15_ carrying CGs, where CG307 and CG15 have played key roles. Yersinabactin and ESBL-encoding mobile genetic elements are uncommon in clinical isolates of *K. variicola* ssp. *variicola* compared to *K. pneumoniae sensu stricto*. Susceptible *K. variicola* ssp. *variicola*, however, is a significant pathogen causing one out of four cases of KpSC bacteraemia in Norway.

## Acknowledgements

We thank The Norwegian Surveillance Program on Resistant Microbes (NORM) for making data available for this study.

## Transparency

None to declare

## Contributions

Study conceivement: A.F, Ø.S, A.S, G.S.S, I.H.L; whole genome sequencing: R.B and E.B.; data analysis: A.F and M.A.K.H; manuscript: A.F and M.A.K.H. All authors contributed to data interpretation, read and commented on the manuscript. Collaborators provided isolates and commented on the final manuscript.

---

**Supplementary figure S1.**
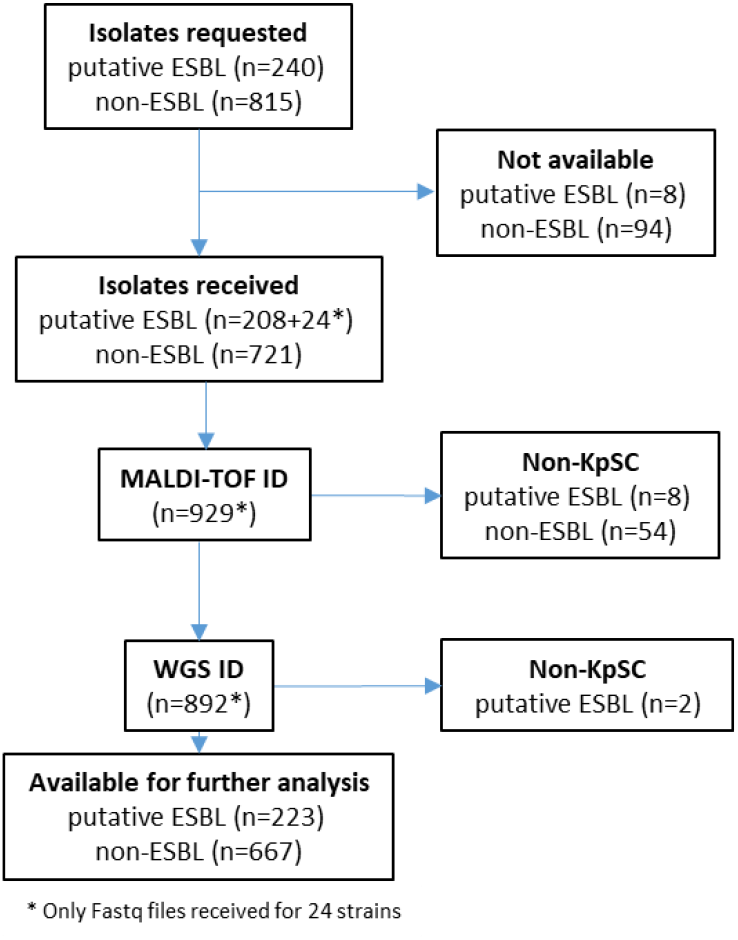
Inclusion of isolates.

**Supplementary figure S2.**
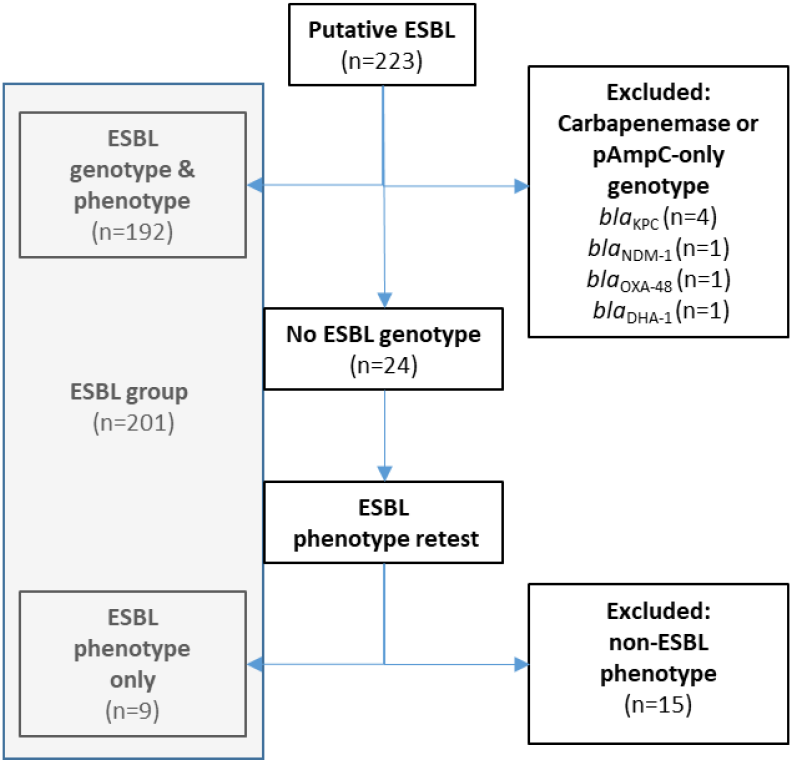
Verification of putative ESBLs.

**Supplementary figure S3.**
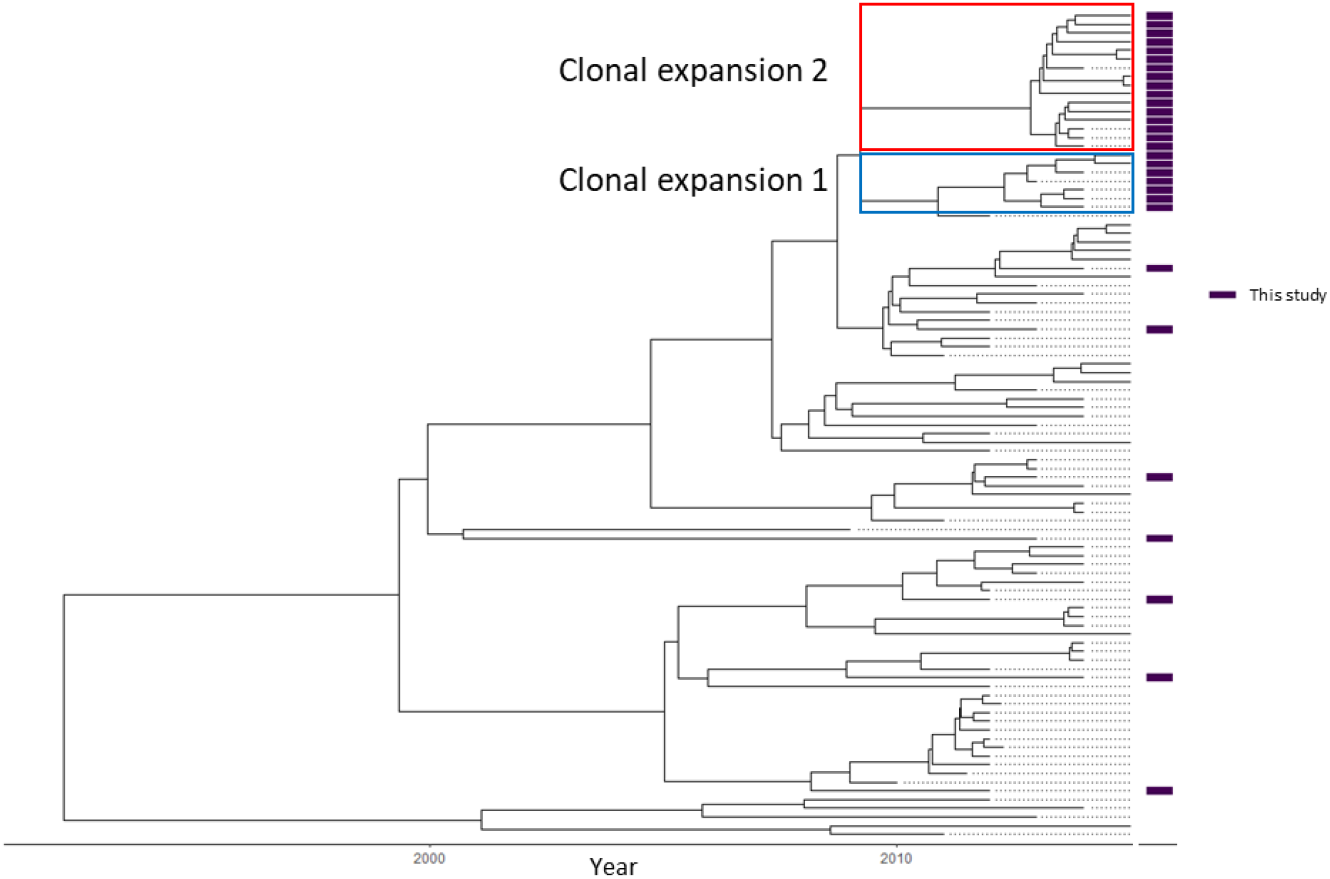
Bayesian phylogeny of global CG307. **Figure adapted from Wyres *et al.* 2019. Isolates in this study marked with purple bars. Proposed clonal expansion 1 and 2 marked with red and blue box, respectively.**

**Supplementary table S2.** Distribution of K-loci amonc clonal groups.

| Supplementary table S2 - Distribution of K-loci among clonal groups |  |  |  |  |
| --- | --- | --- | --- | --- |
| class | K-locus | n | % | Clonal groups (CGs) |
| esbl | unknown | 53 | 26,4 % | CG307 (n=9), CG15 (n=5), CG20 (n=4), CG2947 (n=4), CG258 (n=4), CG35 (n=3), CG70 (n=3), CG45 (n=2), CG1393 (n=1), CG25 (n=1), CG251 (n=1), CG26 (n=1), CG268 (n=1), CG27 (n=1), CG29 (n=1), CG2950 (n=1), CG2952 (n=1), CG37 (n=1), CG377 (n=1), CG39 (n=1), CG392 (n=1), CG405 (n=1), CG48 (n=1), CG551 (n=1), CG661 (n=1), CG716 (n=1), CG76 (n=1) |
| esbl | KL102 | 27 | 13,4 % | CG307 (n=25), CG268 (n=2) |
| esbl | KL2 | 11 | 5,5 % | CG15 (n=9), CG35 (n=1), CG416 (n=1) |
| esbl | KL24 | 11 | 5,5 % | CG45 (n=6), CG15 (n=5) |
| esbl | KL136 | 10 | 5,0 % | CG70 (n=10) |
| esbl | KL38 | 9 | 4,5 % | CG261 (n=5), CG37 (n=3), CG873 (n=1) |
| esbl | KL62 | 9 | 4,5 % | CG48 (n=6), CG198 (n=1), CG2552 (n=1), CG45 (n=1) |
| esbl | KL15 | 8 | 4,0 % | CG258 (n=5), CG37 (n=2), CG392 (n=1) |
| esbl | KL10 | 4 | 2,0 % | CG551 (n=2), CG107 (n=1), CG392 (n=1) |
| esbl | KL17 | 4 | 2,0 % | CG101 (n=3), CG322 (n=1) |
| esbl | KL28 | 4 | 2,0 % | CG20 (n=1), CG26 (n=1), CG2948 (n=1), CG661 (n=1) |
| esbl | KL3 | 4 | 2,0 % | CG896 (n=3), CG873 (n=1) |
| esbl | KL43 | 4 | 2,0 % | CG904 (n=4) |
| esbl | KL51 | 4 | 2,0 % | CG2947 (n=3), CG20 (n=1) |
| esbl | KL112 | 3 | 1,5 % | CG15 (n=3) |
| esbl | KL25 | 3 | 1,5 % | CG20 (n=2), CG45 (n=1) |
| esbl | KL27 | 3 | 1,5 % | CG392 (n=3) |
| esbl | KL48 | 3 | 1,5 % | CG15 (n=3) |
| esbl | KL105 | 2 | 1,0 % | CG1296 (n=1), CG258 (n=1) |
| esbl | KL107 | 2 | 1,0 % | CG377 (n=1), CG659 (n=1) |
| esbl | KL21 | 2 | 1,0 % | CG290 (n=1), CG323 (n=1) |
| esbl | KL46 | 2 | 1,0 % | CG2428 (n=1), CG277 (n=1) |
| esbl | KL5 | 2 | 1,0 % | CG686 (n=2) |
| esbl | KL9 | 2 | 1,0 % | CG22 (n=1), CG719 (n=1) |
| esbl | KL103 | 1 | 0,5 % | CG334 (n=1) |
| esbl | KL108 | 1 | 0,5 % | CG2947 (n=1) |
| esbl | KL13 | 1 | 0,5 % | CG485 (n=1) |
| esbl | KL14 | 1 | 0,5 % | CG37 (n=1) |
| esbl | KL151 | 1 | 0,5 % | CG405 (n=1) |
| esbl | KL19 | 1 | 0,5 % | CG2946 (n=1) |
| esbl | KL20 | 1 | 0,5 % | CG133 (n=1) |
| esbl | KL23 | 1 | 0,5 % | CG584 (n=1) |
| esbl | KL31 | 1 | 0,5 % | CG4246 (n=1) |
| esbl | KL45 | 1 | 0,5 % | CG874 (n=1) |
| esbl | KL53 | 1 | 0,5 % | CG489 (n=1) |
| esbl | KL54 | 1 | 0,5 % | CG29 (n=1) |
| esbl | KL63 | 1 | 0,5 % | CG111 (n=1) |
| esbl | KL64 | 1 | 0,5 % | CG392 (n=1) |
| esbl | KL7 | 1 | 0,5 % | CG4 (n=1) |

| class | K-locus | n | % | Clonal groups (CGs) |
| --- | --- | --- | --- | --- |
| non-esbl | unknown | 305 | 45,7 % | CG26 (n=25), CG20 (n=15), CG37 (n=15), CG35 (n=7), CG641 (n=7), CG643 (n=7), CG10 (n=6), CG1562 (n=6), CG359 (n=6), CG4 (n=5), CG925 (n=5), CG1875 (n=4), CG2947 (n=4), CG3084 (n=4), CG461 (n=4), CG105 (n=3), CG240 (n=3), CG25 (n=3), CG251 (n=3), CG268 (n=3), CG27 (n=3), CG29 (n=3), CG4230 (n=3), CG45 (n=3), CG596 (n=3), CG639 (n=3), CG896 (n=3), CG1114 (n=2), CG1142 (n=2), CG1308 (n=2), CG133 (n=2), CG15 (n=2), CG1423 (n=2), CG1456 (n=2), CG1681 (n=2), CG1777 (n=2), CG197 (n=2), CG200 (n=2), CG250 (n=2), CG338 (n=2), CG347 (n=2), CG3919 (n=2), CG416 (n=2), CG4206 (n=2), CG475 (n=2), CG681 (n=2), CG70 (n=2), CG919 (n=2), CG1035 (n=1), CG1096 (n=1), CG111 (n=1), CG1189 (n=1), CG1193 (n=1), CG1317 (n=1), CG134 (n=1), CG1393 (n=1), CG1485 (n=1), CG1609 (n=1), CG1646 (n=1), CG1664 (n=1), CG1727 (n=1), CG1778 (n=1), CG1791 (n=1), CG180 (n=1), CG1845 (n=1), CG188 (n=1), CG189 (n=1), CG1966 (n=1), CG198 (n=1), CG1980 (n=1), CG2010 (n=1), CG209 (n=1), CG215 (n=1), CG23 (n=1), CG2309 (n=1), CG2386 (n=1), CG2421 (n=1), CG2441 (n=1), CG2445 (n=1), CG2459 (n=1), CG2558 (n=1), CG285 (n=1), CG2850 (n=1), CG292 (n=1), CG294 (n=1), CG2942 (n=1), CG2952 (n=1), CG30 (n=1), CG3048 (n=1), CG3083 (n=1), CG3175 (n=1), CG321 (n=1), CG334 (n=1), CG258 (n=1), CG3506 (n=1), CG3565 (n=1), CG363 (n=1), CG378 (n=1), CG384 (n=1), CG392 (n=1), CG4024 (n=1), CG405 (n=1), CG4076 (n=1), CG4148 (n=1), CG4151 (n=1), CG4156 (n=1), CG4157 (n=1), CG4161 (n=1), CG4163 (n=1), CG4164 (n=1), CG4171 (n=1), CG4174 (n=1), CG4177 (n=1), CG4187 (n=1), CG4189 (n=1), CG4199 (n=1), CG4214 (n=1), CG4218 (n=1), CG4223 (n=1), CG4229 (n=1), CG4243 (n=1), CG4249 (n=1), CG4261 (n=1), CG4281 (n=1), CG4328 (n=1), CG4331 (n=1), CG4333 (n=1), CG4340 (n=1), CG4348 (n=1), CG4359 (n=1), CG4360 (n=1), CG4374 (n=1), CG4376 (n=1), CG4378 (n=1), CG4387 (n=1), CG4389 (n=1), CG4393 (n=1), CG4398 (n=1), CG4400 (n=1), CG477 (n=1), CG48 (n=1), CG485 (n=1), CG499 (n=1), CG501 (n=1), CG515 (n=1), CG584 (n=1), CG616 (n=1), CG628 (n=1), CG686 (n=1), CG697 (n=1), CG76 (n=1), CG767 (n=1), CG86 (n=1), CG889 (n=1), CG942 (n=1) |
| non-esbl | KL28 | 18 | 2,7 % | CG20 (n=11), CG26 (n=4), CG15 (n=1), CG2478 (n=1), CG70 (n=1) |
| non-esbl | KL10 | 17 | 2,5 % | CG359 (n=6), CG26 (n=5), CG198 (n=1), CG252 (n=1), CG515 (n=1), CG551 (n=1), CG584 (n=1), CG76 (n=1) |
| non-esbl | KL2 | 14 | 2,1 % | CG25 (n=7), CG15 (n=4), CG86 (n=2), CG20 (n=1) |
| non-esbl | KL38 | 14 | 2,1 % | CG1423 (n=3), CG20 (n=3), CG37 (n=2), CG641 (n=2), CG3147 (n=1), CG318 (n=1), CG258 (n=1), CG584 (n=1) |
| non-esbl | KL54 | 12 | 1,8 % | CG29 (n=6), CG1778 (n=2), CG3231 (n=1), CG4181 (n=1), CG616 (n=1), CG889 (n=1) |
| non-esbl | KL1 | 11 | 1,6 % | CG23 (n=6), CG2159 (n=1), CG249 (n=1), CG4071 (n=1), CG417 (n=1), CG440 (n=1) |
| non-esbl | KL25 | 11 | 1,6 % | CG134 (n=5), CG20 (n=1), CG27 (n=1), CG2850 (n=1), CG4265 (n=1), CG461 (n=1), CG643 (n=1) |
| non-esbl | KL102 | 10 | 1,5 % | CG10 (n=8), CG105 (n=2) |
| non-esbl | KL24 | 10 | 1,5 % | CG15 (n=4), CG359 (n=2), CG45 (n=2), CG20 (n=1), CG27 (n=1) |
| non-esbl | KL31 | 10 | 1,5 % | CG26 (n=2), CG104 (n=1), CG1096 (n=1), CG188 (n=1), CG1984 (n=1), CG2010 (n=1), CG2108 (n=1), CG4192 (n=1), CG4251 (n=1) |
| non-esbl | KL62 | 10 | 1,5 % | CG461 (n=5), CG643 (n=2), CG160 (n=1), CG45 (n=1), CG664 (n=1) |
| non-esbl | KL21 | 9 | 1,3 % | CG37 (n=2), CG1630 (n=1), CG200 (n=1), CG2609 (n=1), CG290 (n=1), CG323 (n=1), CG4217 (n=1), CG499 (n=1) |
| non-esbl | KL30 | 9 | 1,3 % | CG234 (n=2), CG29 (n=2), CG416 (n=2), CG198 (n=1), CG252 (n=1), CG4239 (n=1) |
| non-esbl | KL63 | 9 | 1,3 % | CG111 (n=3), CG3067 (n=2), CG4203 (n=2), CG4225 (n=1), CG4396 (n=1) |
| non-esbl | KL105 | 8 | 1,2 % | CG1114 (n=6), CG639 (n=1), CG774 (n=1) |
| non-esbl | KL60 | 7 | 1,0 % | CG641 (n=3), CG1224 (n=1), CG1915 (n=1), CG3702 (n=1), CG3919 (n=1) |
| non-esbl | KL103 | 6 | 0,9 % | CG107 (n=1), CG1984 (n=1), CG2010 (n=1), CG3975 (n=1), CG4188 (n=1), CG4352 (n=1) |
| non-esbl | KL134 | 6 | 0,9 % | CG515 (n=5), CG4193 (n=1) |
| non-esbl | KL17 | 6 | 0,9 % | CG322 (n=4), CG101 (n=1), CG2947 (n=1) |
| non-esbl | KL3 | 6 | 0,9 % | CG76 (n=2), CG209 (n=1), CG321 (n=1), CG4192 (n=1), CG4203 (n=1) |
| non-esbl | KL5 | 6 | 0,9 % | CG2421 (n=1), CG2559 (n=1), CG290 (n=1), CG4252 (n=1), CG45 (n=1), CG60 (n=1) |
| non-esbl | KL53 | 6 | 0,9 % | CG1875 (n=3), CG363 (n=2), CG475 (n=1) |
| non-esbl | KL7 | 6 | 0,9 % | CG187 (n=2), CG26 (n=2), CG4 (n=2) |
| non-esbl | KL9 | 6 | 0,9 % | CG197 (n=2), CG1317 (n=1), CG22 (n=1), CG584 (n=1), CG628 (n=1) |
| non-esbl | KL22 | 5 | 0,7 % | CG35 (n=2), CG1193 (n=1), CG248 (n=1), CG3639 (n=1) |
| non-esbl | KL23 | 5 | 0,7 % | CG37 (n=2), CG1262 (n=1), CG20 (n=1), CG4176 (n=1) |
| non-esbl | KL27 | 5 | 0,7 % | CG268 (n=2), CG661 (n=2), CG26 (n=1) |
| non-esbl | KL117 | 4 | 0,6 % | CG2947 (n=2), CG4238 (n=1), CG4290 (n=1) |
| non-esbl | KL13 | 4 | 0,6 % | CG189 (n=1), CG251 (n=1), CG33 (n=1), CG4 (n=1) |
| non-esbl | KL14 | 4 | 0,6 % | CG2441 (n=2), CG363 (n=1), CG37 (n=1) |
| non-esbl | KL143 | 4 | 0,6 % | CG214 (n=3), CG4223 (n=1) |
| non-esbl | KL16 | 4 | 0,6 % | CG15 (n=2), CG215 (n=1), CG4203 (n=1) |
| non-esbl | KL20 | 4 | 0,6 % | CG268 (n=1), CG35 (n=1), CG485 (n=1), CG893 (n=1) |
| non-esbl | KL34 | 4 | 0,6 % | CG1791 (n=1), CG312 (n=1), CG347 (n=1), CG697 (n=1) |
| non-esbl | KL39 | 4 | 0,6 % | CG26 (n=1), CG3370 (n=1), CG3878 (n=1), CG4174 (n=1) |
| non-esbl | KL55 | 4 | 0,6 % | CG20 (n=1), CG392 (n=1), CG4382 (n=1), CG475 (n=1) |
| non-esbl | KL57 | 4 | 0,6 % | CG1035 (n=1), CG3955 (n=1), CG4224 (n=1), CG643 (n=1) |
| non-esbl | KL61 | 4 | 0,6 % | CG1845 (n=1), CG240 (n=1), CG4079 (n=1), CG4228 (n=1) |
| non-esbl | KL114 | 3 | 0,4 % | CG1535 (n=1), CG628 (n=1), CG719 (n=1) |
| non-esbl | KL18 | 3 | 0,4 % | CG101 (n=1), CG4384 (n=1), CG967 (n=1) |
| non-esbl | KL35 | 3 | 0,4 % | CG1096 (n=1), CG4154 (n=1), CG4206 (n=1) |
| non-esbl | KL46 | 3 | 0,4 % | CG2428 (n=1), CG277 (n=1), CG4233 (n=1) |
| non-esbl | KL107 | 2 | 0,3 % | CG4174 (n=1), CG4316 (n=1) |
| non-esbl | KL108 | 2 | 0,3 % | CG129 (n=1), CG461 (n=1) |
| non-esbl | KL11 | 2 | 0,3 % | CG4205 (n=1), CG4349 (n=1) |
| non-esbl | KL110 | 2 | 0,3 % | CG268 (n=1), CG258 (n=1) |
| non-esbl | KL112 | 2 | 0,3 % | CG15 (n=1), CG240 (n=1) |
| non-esbl | KL113 | 2 | 0,3 % | CG184 (n=1), CG925 (n=1) |
| non-esbl | KL116 | 2 | 0,3 % | CG133 (n=2) |
| non-esbl | KL12 | 2 | 0,3 % | CG200 (n=1), CG37 (n=1) |
| non-esbl | KL122 | 2 | 0,3 % | CG10 (n=1), CG70 (n=1) |
| non-esbl | KL124 | 2 | 0,3 % | CG4349 (n=1), CG48 (n=1) |
| non-esbl | KL128 | 2 | 0,3 % | CG189 (n=1), CG4035 (n=1) |
| non-esbl | KL136 | 2 | 0,3 % | CG37 (n=1), CG515 (n=1) |
| non-esbl | KL142 | 2 | 0,3 % | CG26 (n=2) |
| non-esbl | KL149 | 2 | 0,3 % | CG105 (n=1), CG2738 (n=1) |
| non-esbl | KL158 | 2 | 0,3 % | CG397 (n=1), CG776 (n=1) |
| non-esbl | KL48 | 2 | 0,3 % | CG4206 (n=1), CG686 (n=1) |
| non-esbl | KL52 | 2 | 0,3 % | CG501 (n=1), CG873 (n=1) |
| non-esbl | KL64 | 2 | 0,3 % | CG30 (n=1), CG4395 (n=1) |
| non-esbl | KL67 | 2 | 0,3 % | CG1562 (n=1), CG4368 (n=1) |
| non-esbl | KL101 | 1 | 0,1 % | CG767 (n=1) |
| non-esbl | KL111 | 1 | 0,1 % | CG1777 (n=1) |
| non-esbl | KL118 | 1 | 0,1 % | CG322 (n=1) |
| non-esbl | KL120 | 1 | 0,1 % | CG1407 (n=1) |
| non-esbl | KL121 | 1 | 0,1 % | CG107 (n=1) |
| non-esbl | KL123 | 1 | 0,1 % | CG200 (n=1) |
| non-esbl | KL125 | 1 | 0,1 % | CG919 (n=1) |
| non-esbl | KL127 | 1 | 0,1 % | CG440 (n=1) |
| non-esbl | KL130 | 1 | 0,1 % | CG1562 (n=1) |
| non-esbl | KL137 | 1 | 0,1 % | CG4371 (n=1) |
| non-esbl | KL145 | 1 | 0,1 % | CG70 (n=1) |
| non-esbl | KL147 | 1 | 0,1 % | CG20 (n=1) |
| non-esbl | KL162 | 1 | 0,1 % | CG248 (n=1) |
| non-esbl | KL166 | 1 | 0,1 % | CG719 (n=1) |
| non-esbl | KL19 | 1 | 0,1 % | CG1562 (n=1) |
| non-esbl | KL33 | 1 | 0,1 % | CG499 (n=1) |
| non-esbl | KL45 | 1 | 0,1 % | CG4234 (n=1) |
| non-esbl | KL47 | 1 | 0,1 % | CG1562 (n=1) |
| non-esbl | KL49 | 1 | 0,1 % | CG1142 (n=1) |
| non-esbl | KL56 | 1 | 0,1 % | CG475 (n=1) |
| non-esbl | KL58 | 1 | 0,1 % | CG3878 (n=1) |
| non-esbl | KL6 | 1 | 0,1 % | CG4271 (n=1) |
| non-esbl | KL71 | 1 | 0,1 % | CG3938 (n=1) |
| non-esbl | KL74 | 1 | 0,1 % | CG392 (n=1) |
| non-esbl | KL8 | 1 | 0,1 % | CG453 (n=1) |
| non-esbl | KL81 | 1 | 0,1 % | CG252 (n=1) |

**Supplementary table S3.**
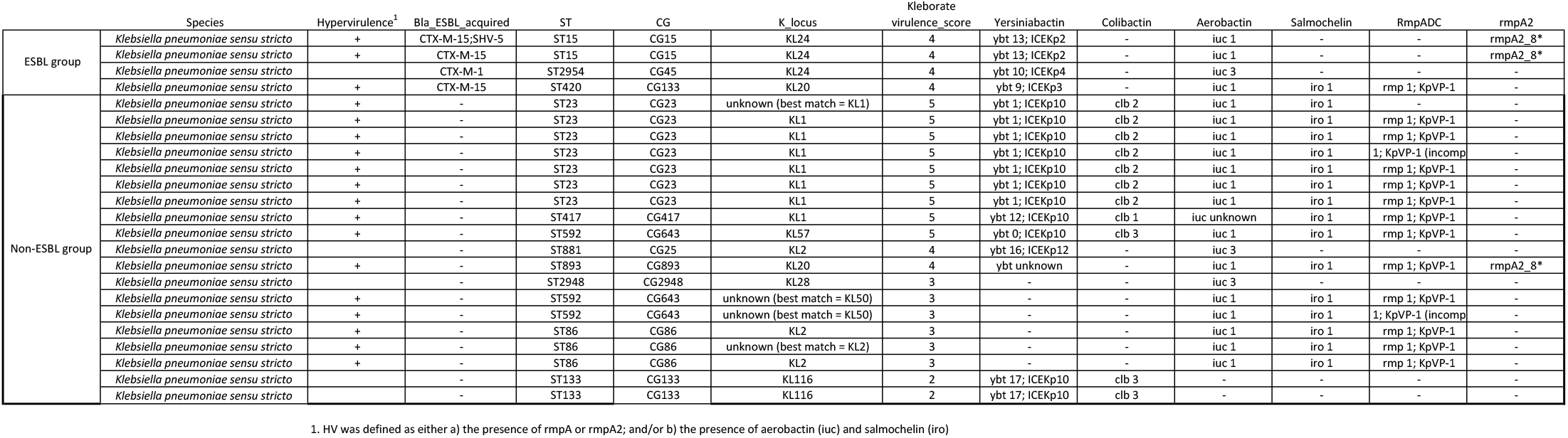
Distribution of virulence loci.

**Supplementary table S4.** Replicon distribution.

|  |  | Replicon family |  |  |  |  |  |  |  |  |  |  |  |  |  |  |  |  |  |  |  |
| --- | --- | --- | --- | --- | --- | --- | --- | --- | --- | --- | --- | --- | --- | --- | --- | --- | --- | --- | --- | --- | --- |
|  | species | n | Col | IncA | IncB | IncC | IncFIA | IncFIB | IncFII | IncH | IncI | IncL | IncM | IncN | IncQ | IncR | IncU | IncX | IncY | rep_a | pKP1433 |
| ESBL | total | 201 | 62.7 % (n=126) | 2.0 % (n=4) | 1.5 % (n=3) | 2.0 % (n=4) | 6.0 % (n=12) | 87.6 % (n=176) | 82.6 % (n=166) | 6.5 % (n=13) | 1.5 % (n=3) | 1.5 % (n=3) | 1.5 % (n=3) | 4.5 % (n=9) | 6.0 % (n=12) | 18.9 % (n=38) |  | 2.0 % (n=4) | 0.5 % (n=1) |  |  |
|  | *Klebsiella pneumoniae* | 198 | 63.1 % (n=125) | 2.0 % (n=4) | 1.5 % (n=3) | 2.0 % (n=4) | 6.1 % (n=12) | 87.9 % (n=174) | 82.8 % (n=164) | 6.1 % (n=12) | 1.5 % (n=3) | 1.0 % (n=2) | 1.0 % (n=2) | 4.5 % (n=9) | 6.1 % (n=12) | 18.2 % (n=36) |  | 2.0 % (n=4) | 0.5 % (n=1) |  |  |
|  | *Klebsiella quasipneumoniae* ssp. *similipneumoniae* | 2 | 50.0 % (n=1) |  |  |  |  | 100.0 % (n=2) | 50.0 % (n=1) | 50.0 % (n=1) |  | 50.0 % (n=1) | 50.0 % (n=1) |  |  | 50.0 % (n=1) |  |  |  |  |  |
|  | *Klebsiella varicola* | 1 |  |  |  |  |  |  | 100.0 % (n=1) |  |  |  |  |  |  | 100.0 % (n=1) |  |  |  |  |  |
| Non-ESBL | total | 667 | 58.9 % (n=393) |  | 0.1 % (n=1) |  | 15.0 % (n=100) | 72.0 % (n=480) | 48.7 % (n=325) | 5.7 % (n=38) | 0.3 % (n=2) | 0.6 % (n=4) | 0.7 % (n=5) | 3.0 % (n=20) | 0.3 % (n=2) | 14.5 % (n=97) | 0.1 % (n=1) |  |  | 0.9 % (n=6) | 0.1 % (n=1) |
|  | *Klebsiella pneumoniae* | 461 | 63.3 % (n=292) |  | 0.2 % (n=1) |  | 14.3 % (n=66) | 79.4 % (n=366) | 54.2 % (n=250) | 7.6 % (n=35) | 0.4 % (n=2) | 0.7 % (n=3) | 0.9 % (n=4) | 3.5 % (n=16) | 0.4 % (n=2) | 14.1 % (n=65) | 0.2 % (n=1) |  |  | 0.9 % (n=4) | 0.2 % (n=1) |
|  | *Klebsiella quasipneumoniae* ssp. *quasipneumoniae* | 21 | 47.6 % (n=10) |  |  |  | 9.5 % (n=2) | 33.3 % (n=7) | 14.3 % (n=3) |  |  | 4.8 % (n=1) | 4.8 % (n=1) | 9.5 % (n=2) |  | 14.3 % (n=3) |  |  |  |  |  |
|  | *Klebsiella quasipneumoniae* ssp. *similipneumoniae* | 22 | 59.1 % (n=13) |  |  |  | 13.6 % (n=3) | 54.5 % (n=12) | 27.3 % (n=6) |  |  |  |  |  |  | 40.9 % (n=9) |  |  |  |  |  |
|  | *Klebsiella varicola* ssp. *varicola* | 163 | 47.9 % (n=78) |  |  |  | 17.8 % (n=29) | 58.3 % (n=95) | 40.5 % (n=66) | 1.8 % (n=3) |  |  |  | 1.2 % (n=2) |  | 12.3 % (n=20) |  |  |  | 1.2 % (n=2) |  |

**Supplementary table S5.** Temporal and geographical distribution, and SNP ranges in prevalent CGs.

| SNPs difference within clonal group |  |  |  |  |  |  |  |  |
| --- | --- | --- | --- | --- | --- | --- | --- | --- |
| Clonal group (CG) | n | min | max | median | IQR | Year first detected | Found at least once during 2001-2015 in regions: | Sequence types(STs) within clonal group |
| CG15 | 39 | 0 | 12439 | 8806 | 9667 | 2001 | East, Middle, North, Oslo_Akershus, South, West | ST14, ST15, ST627 |
| CG20 | 43 | 0 | 14379 | 6930 | 8509 | 2001 | East, Middle, North, Oslo_Akershus, South, West | ST1412, ST16, ST17, ST20, ST22, ST336, ST422, ST4285, ST636 |
| CG26 | 44 | 0 | 20488 | 7564 | 3317.75 | 2001 | East, Middle, North, Oslo_Akershus, South, West | ST163, ST2116, ST2211, ST253, ST26, ST3043, ST3191, ST4273, ST4279, ST4383, ST704 |
| CG70 | 18 | 0 | 14694 | 52.5 | 12829 | 2001 | East, Middle, Oslo_Akershus, South, West | ST1873, ST2344, ST4283, ST70 |
| CG307 | 34 | 0 | 109 | 39 | 43 | 2012 | East, Middle, North, Oslo_Akershus, South, West | ST307 |
| CG48 | 9 | 0 | 21449 | 166 | 2122 | 2001 | East, Middle, Oslo_Akershus, West | ST1411, ST48 |
| CG45 | 17 | 0 | 11319 | 2885 | 3032 | 2005 | East, Middle, Oslo_Akershus, South, West | ST2954, ST45, ST485 |
| CG258 | 13 | 0 | 12451 | 3559 | 7295 | 2009 | East, Middle, North, Oslo_Akershus, South, West | ST11, ST340, ST437 |

## Supplementary methods

### Whole genome sequencing

All study isolates were subject to short-read WGS. DNA was extracted using the MagNAPure 96 system (Roche Applied Science, Manheim, Germany) and sequencing libraries were prepared according to the Nextera XT DNA Library preparation protocol (Illumina, San Diego, CA, USA). WGS was performed on an Illumina MiSeq platform with v3 chemistry to generate 2×300 bp paired-end reads. For isolates where only FASTQ files were received, sequencing had been performed on an Illumina HiSeq 2500 platform at Eurofins Genomics (Eurofins Genomics Europe, Konstanz, Germany) generating 2×125 bp paired-end reads.

To achieve closed genomes for selected isolates, additional long read WGS was performed. DNA was extracted manually using the Beckman Coulter Life Science GenFind V3 kit (C34881) according to the supplemental protocol ‘DNA extraction from Bacteria using GenFind v3’ (Beckman Coulter, Brea, CA, USA). DNA libraries were prepared using the 1D Ligation sequencing kit (SQK-LSK108) and the Native barcoding kit (EXP-NBD103) (Oxford Nanopore Technologies [ONT], Oxford, United Kingdom) according to the ONT protocol ‘native barcoding genomic DNA’ or ‘genomic DNA by ligation’ without shearing to maximize the sequencing read length. Finally, libraries were loaded onto a R9.4.1 MinION flow cell (FLO-MIN106) or a R9.4.1 Flongle flow cell (FLO-FLG001) and sequenced on the ONT MinION Mk1B device (MIN-101B).

### In-silico analyses

Short-read sequences were trimmed based on quality and adapter content with TrimGalore v0.5.0 ^1^ and *de novo* assembly was performed with Unicycler v0.4.8 ^2^, which uses SPAdes v3.13.1 ^3^ for assembly and Pilon v1.23 ^4^ for polishing. Kleborate v2.0.4^5^ was used to identify species and determine of multilocus sequence type (ST), virulence loci and AMR genes (CARD database v3.0.8 ^6^) from assembled genomes. ). Kaptive ^7^ was used to identify capsule (K) biosynthesis loci reporting calls with confidence level “Good” or higher. Putative ESBLs with no definite ESBL-gene had their read sets investigated using SRST2 v.2.0.4^8^ with CARD database v3.0.8, as some reads may have been discarded in the assembly process. Only ESBL-gene matches with 100 % sequence coverage and identity were included in further analyses. Plasmid replicons were identified with SRST2 v0.2.0 using the Plasmidfinder database version 2021-01-13. ^9^

A core chromosomal single-nucleotide variant (SNV) alignment of the verified ESBL and non-ESBL genomes was generated to assess their relatedness. The short-reads were mapped to the chromosome of the ST23 reference genome NTUH-K2044 (NC_01273.1) with the RedDog V1beta.11 ^10^ pipeline, using Bowtie2 v2.3.4.2^11^ for read mapping and SAMtools v1.9^12^ for SNV calling. RedDog was used with default parameters as described previously ^13^, except for the read depth threshold which was set to ≥8 (default ≥10) to include all genomes. A maximum likelihood (ML) phylogeny was inferred from the resulting alignment (868 genomes, 867815 SNPs) using FastTree v2.1.10 (gamma distribution of rate heterogeneity among sites)^14^.

Clonal groups (CGs) were defined by patristic distances. This method was chosen as it has previously been used to cluster CGs in KpSC (Bialek-Davenet, S., et al., 2019) and a distance threshold of 0.04 was used as it grouped STs that have previously been identified as belonging to clinically distinct CGs. The CGs dominated by ST14 and ST340 where denoted CG15 and CG258, as these names are more commonly used. ^15, 16^.

Long-read sequences were base called in barcode-trimmed and de-multiplexed high-accuracy mode using Guppy Basecalling Software v3.2.4+d9ed22f^17,16,17,16^[16][16][16][16][16](16), followed by quality filtering with Filtlong v0.2.0^18^. To resolve the complete genome sequence of these isolates, hybrid assembly with the corresponding short-read genomes using Unicycler v0.4.8 was performed. The completed genomes were subsequently annotated with the NCBI Prokaryotic Annotation Pipeline v5.1 ^19^, using default parameters.

To assess the clonal relatedness of the ST307 genomes, an alignment was generated with RedDog, using the hybrid-assembled closed ST307 genome as the mapping reference . The resulting alignment with 391 variant sites was screened for recombination events using Gubbins v2.3.4 ^20^ with convergence method “weighted Robinson-Foulds”. This alignment was passed to RAxML v8.2.10^21^ to infer ML phylogeny. The best-scoring ML tree was chosen from five independent runs with the GTR+ nucleotide substitution model, followed by a rapid bootstrap analysis (100 replicates) to estimate branch support.

## References

1. Podschun R, Ullmann U. Klebsiella spp. as nosocomial pathogens: epidemiology, taxonomy, typing methods, and pathogenicity factors. Clin Microbiol Rev 1998; 11: 589–603.

2. Navon-Venezia S, Kondratyeva K, Carattoli A. Klebsiella pneumoniae: a major worldwide source and shuttle for antibiotic resistance. FEMS Microbiol Rev 2017; 41: 252–75.

3. Tacconelli E, Carrara E, Savoldi A et al. Discovery, research, and development of new antibiotics: the WHO priority list of antibiotic-resistant bacteria and tuberculosis. The Lancet Infectious diseases 2018; 18: 318–27.

4. Magiorakos AP, Srinivasan A, Carey RB et al. Multidrug-resistant, extensively drug-resistant and pandrug-resistant bacteria: an international expert proposal for interim standard definitions for acquired resistance. Clinical microbiology and infection: the official publication of the European Society of Clinical Microbiology and Infectious Diseases 2012; 18: 268–81.

5. Wyres KL, Holt KE. Klebsiella pneumoniae Population Genomics and Antimicrobial-Resistant Clones. Trends in microbiology 2016; 24: 944–56.

6. Wyres KL, Hawkey J, Hetland MAK et al. Emergence and rapid global dissemination of CTX-M-15-associated Klebsiella pneumoniae strain ST307. The Journal of antimicrobial chemotherapy 2019; 74: 577–81.

7. Villa L, Feudi C, Fortini D et al. Diversity, virulence, and antimicrobial resistance of the KPC-producing Klebsiella pneumoniae ST307 clone. Microbial genomics 2017; 3: e000110.

8. David S, Cohen v, Reuter S et al. Integrated chromosomal and plasmid sequence analyses reveal diverse modes of carbapenemase gene spread among Klebsiella pneumoniae. Proceedings of the National Academy of Sciences of the United States of America 2020.

9. Laxminarayan R, Duse A, Wattal C et al. Antibiotic resistance-the need for global solutions. The Lancet Infectious diseases 2013; 13: 1057–98.

10. Russo TA, Marr CM. Hypervirulent Klebsiella pneumoniae. Clin Microbiol Rev 2019; 32.

11. Lam MMC, Wyres KL, Duchene S et al. Population genomics of hypervirulent Klebsiella pneumoniae clonal-group 23 reveals early emergence and rapid global dissemination. Nature communications 2018; 9: 2703.

12. Bialek-Davenet S, Criscuolo A, Ailloud F et al. Genomic definition of hypervirulent and multidrug-resistant Klebsiella pneumoniae clonal groups. Emerging infectious diseases 2014; 20: 1812–20.

13. Wyres KL, Wick RR, Judd LM et al. Distinct evolutionary dynamics of horizontal gene transfer in drug resistant and virulent clones of Klebsiella pneumoniae. PLoS genetics 2019; 15: e1008114.

14. Rodrigues C, Passet V, Rakotondrasoa A et al. Description of Klebsiella africanensis sp. nov., Klebsiella variicola subsp. tropicalensis subsp. nov. and Klebsiella variicola subsp. variicola subsp. nov. Research in microbiology 2019; 170: 165–70.

15. Rodrigues C, Passet v, Rakotondrasoa A et al. Identification of Klebsiella pneumoniae, Klebsiella quasipneumoniae, Klebsiella variicola and Related Phylogroups by MALDI-TOF Mass Spectrometry. Frontiers in microbiology 2018; 9: 3000.

16. Maatallah M, Vading M, Kabir MH et al. Klebsiella variicola is a frequent cause of bloodstream infection in the stockholm area, and associated with higher mortality compared to K. pneumoniae. PloS one 2014; 9: e113539.

17. EUCAST. Clinical breakpoints - breakpoints and guidance.

18. NORM/NORM-VET 2019. Usage of Antimicrobial Agents and Occurrence of Antimicrobial Resistance in Norway. In: Simonsen GS, Blix HS, Grave K et al., eds, 2020.

19. Tofteland S, Haldorsen B, Dahl KH et al. Effects of phenotype and genotype on methods for detection of extended-spectrum-beta-lactamase-producing clinical isolates of Escherichia coli and Klebsiella pneumoniae in Norway. J Clin Microbiol 2007; 45: 199–205.

20. Wick RR, Judd LM, Gorrie CL et al. Unicycler: Resolving bacterial genome assemblies from short and long sequencing reads. PLoS computational biology 2017; 13: e1005595.

21. Lam MMC, Wick RR, Watts SC et al. A genomic surveillance framework and genotyping tool for Klebsiella pneumoniae and its related species complex. Nature communications 2021; 12: 4188.

22. Alcock BP, Raphenya AR, Lau TTY et al. CARD 2020: antibiotic resistome surveillance with the comprehensive antibiotic resistance database. Nucleic Acids Res 2020; 48: D517–d25.

23. Wyres KL, Wick RR, Gorrie C et al. Identification of Klebsiella capsule synthesis loci from whole genome data. Microbial genomics 2016; 2: e000102.

24. Inouye M, Conway TC, Zobel J et al. Short read sequence typing (SRST): multi-locus sequence types from short reads. BMC Genomics 2012; 13: 338.

25. Carattoli A, Zankari E, Garcia-Fernandez A et al. In silico detection and typing of plasmids using PlasmidFinder and plasmid multilocus sequence typing. Antimicrobial agents and chemotherapy 2014; 58: 3895–903.

26. Holt K. RedDog. https://github.com/katholt/reddog.

27. Price MN, Dehal PS, Arkin AP. FastTree 2--approximately maximum-likelihood trees for large alignments. PloS one 2010; 5: e9490.

28. Valencia-Bacca J, Silva MM, Cerdeira L et al. Detection and Whole-Genome Analysis of a High-Risk Clone of Klebsiella pneumoniae ST340/CG258 Producing CTX-M-15 in a Companion Animal. Microb Drug Resist 2020; 26: 611–5.

29. Stamatakis A. RAxML version 8: a tool for phylogenetic analysis and post-analysis of large phylogenies. Bioinformatics 2014; 30: 1312–3.

30. Huynh BT, Passet V, Rakotondrasoa A et al. Klebsiella pneumoniae carriage in low-income countries: antimicrobial resistance, genomic diversity and risk factors. Gut Microbes 2020; 11: 1287–99.

31. R: A language and environment for statistical computing. In: Team RC, ed: R Foundation for Statistical Computing, Vienna, Austria, 2020.

32. Lam MMC, Wyres KL, Wick RR et al. Convergence of virulence and MDR in a single plasmid vector in MDR Klebsiella pneumoniae ST15. The Journal of antimicrobial chemotherapy 2019; 74: 1218–22.

33. Long SW, Olsen RJ, Eagar TN et al. Population Genomic Analysis of 1,777 Extended-Spectrum Beta-Lactamase-Producing Klebsiella pneumoniae Isolates, Houston, Texas: Unexpected Abundance of Clonal Group 307. mBio 2017; 8.

34. Moradigaravand D, Martin V, Peacock SJ et al. Evolution and Epidemiology of Multidrug-Resistant Klebsiella pneumoniae in the United Kingdom and Ireland. mBio 2017; 8.

35. Gorrie CL, Mirceta M, Wick RR et al. Gastrointestinal Carriage Is a Major Reservoir of Klebsiella pneumoniae Infection in Intensive Care Patients. Clin Infect Dis 2017; 65: 208–15.

36. Martin RM, Cao J, Brisse S et al. Molecular Epidemiology of Colonizing and Infecting Isolates of Klebsiella pneumoniae. mSphere 2016; 1.

37. Raffelsberger N, Hetland MAK, Svendsen K et al. Gastrointestinal carriage of Klebsiella pneumoniae in a general adult population: a cross-sectional study of risk factors and bacterial genomic diversity. Gut Microbes 2021; 13: 1939599.

38. ECDC. European Centre for Disease Prevention and Control. Antimicrobial resistance surveillance in Europe 2015. Annual Report of the European Antimicrobial Resistance Surveillance Network (EARS-Net). Stockholm, 2017.

39. Lam MMC, Wick RR, Wyres KL et al. Genetic diversity, mobilisation and spread of the yersiniabactin-encoding mobile element ICEKp in Klebsiella pneumoniae populations. Microbial genomics 2018; 4.

40. Carattoli A. Plasmids and the spread of resistance. International journal of medical microbiology: IJMM 2013; 303: 298–304.

## References

1. Krueger F. TrimGalore. https://github.com/FelixKrueger/TrimGalore.

2. Wick RR, Judd LM, Gorrie CL et al. Unicycler: Resolving bacterial genome assemblies from short and long sequencing reads. PLoS computational biology 2017; 13: e1005595.

3. Bankevich A, Nurk S, Antipov D et al. SPAdes: a new genome assembly algorithm and its applications to single-cell sequencing. Journal of computational biology: a journal of computational molecular cell biology 2012; 19: 455–77.

4. Walker BJ, Abeel T, Shea T et al. Pilon: an integrated tool for comprehensive microbial variant detection and genome assembly improvement. PloS one 2014; 9: e112963.

5. Lam MMC, Wick RR, Watts SC et al. A genomic surveillance framework and genotyping tool for Klebsiella pneumoniae and its related species complex. Nature communications 2021; 12: 4188.

6. Alcock BP, Raphenya AR, Lau TTY et al. CARD 2020: antibiotic resistome surveillance with the comprehensive antibiotic resistance database. Nucleic Acids Res 2020; 48: D517–d25.

7. Wyres KL, Wick RR, Gorrie C et al. Identification of Klebsiella capsule synthesis loci from whole genome data. Microbial genomics 2016; 2: e000102.

8. Inouye M, Conway TC, Zobel J et al. Short read sequence typing (SRST): multi-locus sequence types from short reads. BMC Genomics 2012; 13: 338.

9. Carattoli A, Zankari E, Garcia-Fernandez A et al. In silico detection and typing of plasmids using PlasmidFinder and plasmid multilocus sequence typing. Antimicrobial agents and chemotherapy 2014; 58: 3895–903.

10. Holt K. RedDog. https://github.com/katholt/reddog.

11. Langmead B, Salzberg SL. Fast gapped-read alignment with Bowtie 2. Nat Methods 2012; 9: 357–9.

12. Li H, Handsaker B, Wysoker A et al. The Sequence Alignment/Map format and SAMtools. Bioinformatics 2009; 25: 2078–9.

14. Price MN, Dehal PS, Arkin AP. FastTree 2--approximately maximum-likelihood trees for large alignments. PloS one 2010; 5: e9490.

15. Valencia-Bacca J, Silva MM, Cerdeira L et al. Detection and Whole-Genome Analysis of a High-Risk Clone of Klebsiella pneumoniae ST340/CG258 Producing CTX-M-15 in a Companion Animal. Microb Drug Resist 2020; 26: 611–5.

16. Navon-Venezia S, Kondratyeva K, Carattoli A. Klebsiella pneumoniae: a major worldwide source and shuttle for antibiotic resistance. FEMS Microbiol Rev 2017; 41: 252–75.

17. Guppy. https://community.nanoporetech.com.

18. Wick RR. Filtlong. https://github.com/rrwick/Filtlong.

19. Tatusova T, DiCuccio M, Badretdin A et al. NCBI prokaryotic genome annotation pipeline. Nucleic Acids Res 2016; 44: 6614–24.

20. Croucher NJ, Page AJ, Connor TR et al. Rapid phylogenetic analysis of large samples of recombinant bacterial whole genome sequences using Gubbins. Nucleic Acids Res 2015; 43: e15.

21. Stamatakis A. RAxML version 8: a tool for phylogenetic analysis and post-analysis of large phylogenies. Bioinformatics 2014; 30: 1312–3.

